# TOP the Transcription Orientation Pipeline and its use to investigate the transcription of non-coding regions: assessment with CRISPR direct repeats and intergenic sequences

**DOI:** 10.1101/2020.01.15.903914

**Authors:** Kimberley Houenoussi, Roudaina Boukheloua, Jean-Philippe Vernadet, Daniel Gautheret, Gilles Vergnaud, Christine Pourcel

## Abstract

A large proportion of non-coding sequences in prokaryotes are transcribed, playing an important role in the cell metabolism and defense against exogenous elements. This is the case of small RNAs and of clustered regularly interspaced short palindromic repeats “CRISPR” arrays. The CRISPR-Cas system is a defense mechanism that protects bacterial and archaeal genomes against invasions by mobile genetic elements such as viruses and plasmids. The CRISPR array, made of repeats separated by unique sequences called spacers, is transcribed but the nature of the promoter and of the transcription regulation is not well known. We describe the Transcription Orientation Pipeline (TOP) which makes use of transcriptome sequence reads to recover those corresponding to a selected sequence, and determine the direction of the transcription. CRISPR repeat sequences extracted from CRISPRCasdb were used to test the performances of the program. Statistical tests show that CRISPR elements can be reliably oriented with as little as 100 mapped reads. TOP was applied to all the available RNA-Seq Illumina sequencing archives from species possessing a CRISPR array, allowing comparisons with programs dedicated to the orientation of CRISPR repeats. In addition TOP was used to analyze small non-coding RNAs in *Staphylococcus aureus*, demonstrating that it is a valuable and convenient tool to investigate the transcription orientation of any sequence of interest.

**Availability and implementation:** TOPs is implemented in Python and is freely available via the I2BC github repository at https://github.com/i2bc/TOP.

## INTRODUCTION

Small RNAs (sRNAs), a category of noncoding RNAs (ncRNAs) different from rRNAs and tRNAs, are abundant and diverse in Archaea and Bacteria (Gomes-Filho *et al.*, 2018; Wagner & Romby, 2015). They are known to be vital actors in the regulation of gene expression and for some of them in mechanisms of defense against exogenous DNA. A particular case is that of CRISPR arrays, remarkable structures known to interact with a set of proteins called Cas (CRISPR Associated) (Haft *et al.*, 2005; Horvath & Barrangou, 2010) to form the adaptive CRISPR-Cas immune systems. Functional analysis of CRISPR-Cas systems in different species showed that they can play a number of roles including defense against foreign genetic elements such as virus and plasmids, regulation of lysogeny, and regulation of biofilm formation (Hille *et al.*, 2018). A CRISPR locus is made of a succession of repeats of 23 to 50 bp in length interspaced by similarly sized unique regions called spacers. Some CRISPR arrays are preceded by a 100 – 500 bp promoter region also called the “leader”. Spacers provide the specificity of the defense mechanism and mostly originate from phages or plasmids (Pourcel *et al.*, 2005). *Cas* genes are found in most CRISPR containing genomes. Depending on the set of Cas, six types and 33 subtypes are distinguished, among which significant sequence conservation has been demonstrated (Koonin *et al.*, 2017; Koonin & Makarova, 2019).

Several bioinformatics tools have been developed in order to study CRISPR arrays but identification of the promoter for the CRISPR transcription is not always straightforward (Alkhnbashi *et al.*, 2019). Tools were developed based on characteristics of the arrays to predict the repeat orientation (CRISPRdirection (Biswas *et al.*, 2014) CRISPRstrand (Alkhnbashi *et al.*, 2014) CRISPRmap (Lange *et al.*, 2013)) as well as the presence of a leader (CRISPRleader) (Alkhnbashi *et al.*, 2016).

Here we have used transcriptomic data to identify the orientation of the CRISPR transcription and hence the repeat direction. Previously the work of Heidrich et al. (Heidrich *et al.*, 2015) and Ye and Zhang (Ye & Zhang, 2016) showed that this approach was very informative to investigate CRISPR-Cas system expression. We developed a pipeline that searches for transcripts corresponding to CRISPR repeats present in bacteria and archaea, and then determines the orientation of the repeat. As a control we analyzed repeats of CRISPR arrays which transcription was experimentally determined, as well as other small non-coding RNAs from *Staphylococcus aureus*. We show that this approach allows a fast investigation of CRISPR transcription and improvement of orientation tools.

## METHODS

### Environment and Input

The Transcription Orientation Pipeline “TOP” was written in Python 3 and Snakemake and packaged with conda.

A list of direct repeat sequences was downloaded in text format from CRISPRCasdb (Pourcel *et al.*, 2019) and labelled by the taxonomy identifier (taxid) of the species of origin. Only repeats of CRISPR arrays with the highest evidence level (evidence level 4 or el4) as determined by CRISPRCasFinder were retained (Couvin *et al.*, 2018). The transcriptome RNA-Seq archives corresponding to the same taxid were recovered from the “European Nucleotide Archive” (ENA) database (https://www.ebi.ac.uk/ena).

The archives were stored into a folder that carries the name of the taxid, then used as input into the pipeline.

### Programs used in the pipeline

BBDuk (https://sourceforge.net/projects/bbmap/) is a sequence reads decontamination tool based on k-mers search that can extract reads containing any given sequence or its reverse complement. BBDuk was used to extract reads matching an input sequence. rnaSPAdes (https://github.com/ablab/spades), was used for *de novo* transcriptome assembly of the extracted RNA-Seq reads (Bankevich *et al.*, 2012; Bushmanova *et al.*, 2019). BWA (Li & Durbin, 2009) was used to map the RNA-Seq reads on contigs. Seqtk (https://github.com/lh3/seqtk) was used to extract a subset of reads from a fastqfile. RSeQC (Wang *et al.*, 2012) available at http://rseqc.sourceforge.net/ uses an annotation file in .bed format and an alignment file (SAM/BAM). We used it to calculate the percentage of reads mapped in the sense (+) or antisense (-) orientation relative to annotated genes.

Geneious Prime2019 (Biomatters, New-Zealand) was used to produce graphic visualizations of mapped reads.

### Archive orientation

Using Seqtk and fastq files of archives as inputs, 500 000 reads are randomly selected and mapped onto the corresponding reference genome with BWA. Read orientation is calculated with RseQC based on the reference genome annotation. If a majority of reads (default is >= 65 %) is mapped on one strand of the genes, the archive is considered as oriented. Archives are labelled as unstranded, + (sense) or – (antisense).

### Repeat orientation

Orientations of the archives and of the repeat-containing reads are compared to generate the orientation of the CRISPR repeats.

### Statistical test

For each CRISPR element in stranded libraries, the number of sense reads (sr) and of total mapped reads (tr) are used to test the H0 hypothesis that the distribution of repeats orientation is not oriented at random, using a binomial test: binom.test (sr, tr, p = 0.5). Distributions of p-values and numbers of reads in each archive are plotted using the ggplot R package. Alternatively, the log of the number of reads was used to generate graphs.

## RESULTS

### Selecting the archives and evaluating their orientation

Depending on the technique used to prepare the libraries before sequencing, RNA-seq archives may be unstranded or oriented sense or antisense relative to gene transcription. Only orientated archives are of interest for our analyses. To determine the orientation we made use of Seqtk and RSeQC which evaluate, using a portion of the total reads archive, the transcription of selected genes and provide the percentage of reads in sense and antisense orientation as output.

### Design of the pipeline for CRISPR transcription analysis

The first step in the pipeline was the search for reads specific for the input sequence, i.e. the repeat in the case of a CRISPR analysis. We used BBDuk, a k-mer search software to perform this task. BBDuk takes as input a k-mer size and a fastq file: k-mer sizes from 17 to 19 were evaluated. Because CRISPR repeats range in size from 23bp to 50bp, shorter k-mers increase the risk of recovering non-specific reads while longer k-mer reduces sensitivity.

To test the effect of k-mer size, three bacterial genomes were selected harboring from one to 15 CRISPR arrays and in which the orientation of the CRISPR transcription had been experimentally demonstrated: *Pseudomonas aeruginosa* (taxid 287), *Xanthomonas oryzae* (taxid 347) and *Methanocaldococcus jannaschii* (taxid 2190). One stranded ENA archive was randomly selected for each genome. BBDuk was used to extract reads corresponding to the consensus repeat sequence present in the el4 CRISPRs in each of these strains, as found by CRISPRCasFinder (Couvin *et al.*, 2018). In the three assays, a portion of the recovered reads mapped at positions other than CRISPR arrays, as shown for *M. janaschii* using Geneious on Figure 1.

**Figure 1:**
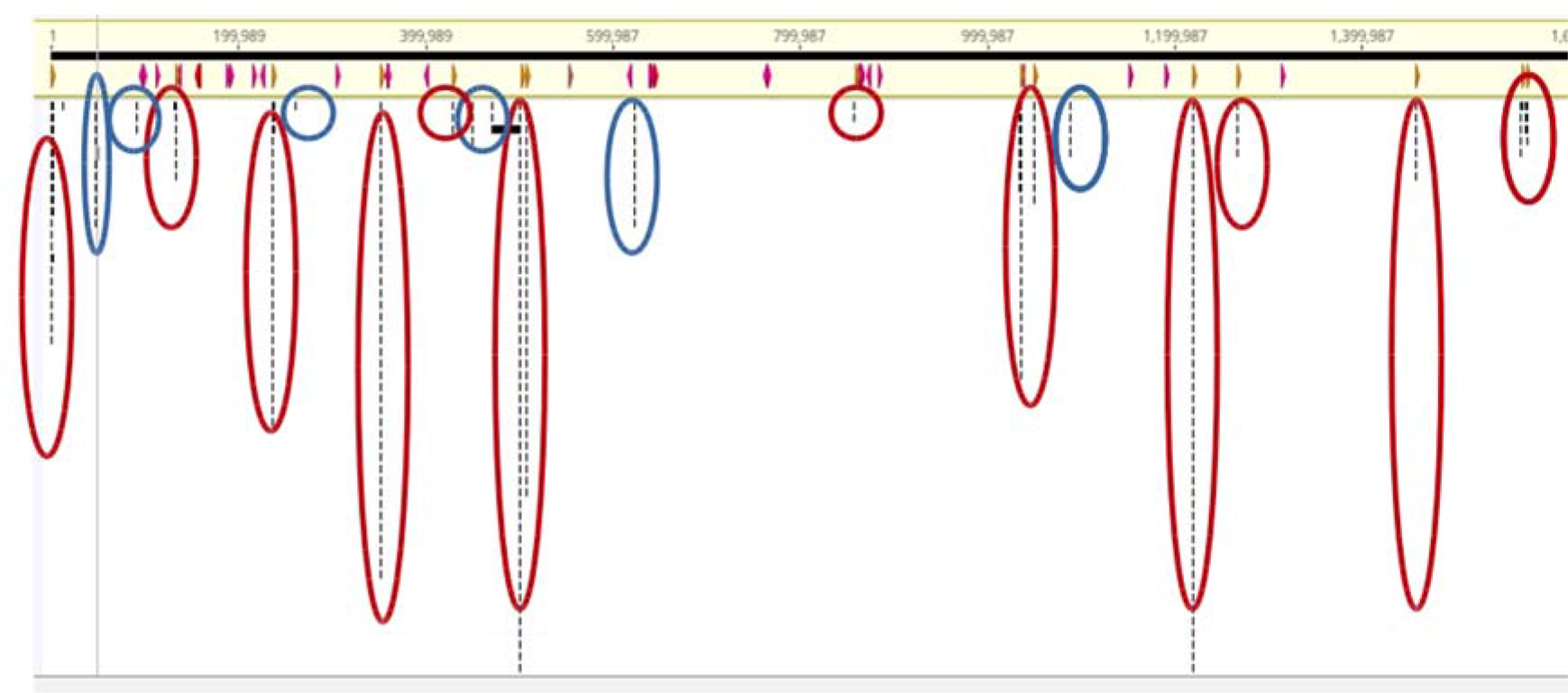
Mapping of RNA-Seq transcriptome reads of *M. jannaschii* on the full genome. A k-mer of size 19 was used to extract reads specific for the CRISPR repeat from *M. jannaschii DSM* 2261 genome in the SRR4240639 archive. Reads recovered by BBDuk were mapped onto the full genome using Geneious prime. In red the CRISPR arrays and in blue other genomic regions indicating that non-specific reads are recovered.

Increasing the size of k did not increase specificity but led to an important loss of reads. From the different tests performed with the three model genomes, we selected a k-mer size of 19 which will be used by default to run the program. This can be modified to suit different types of sequences used as input.

In order to filter out non-specific reads recovered by BBDuk, we assembled the BBDuk output into contigs using rnaSPAdes, retaining only those on which the queried repeat (input sequence) matched perfectly at least once. This assembly step is made necessary by the high polymorphism of CRISPR elements, which means actual elements found in a given ENA library often differ from the corresponding reference genome for this species. In the next step the BBDuk output was mapped onto the assembled contig(s) using BWA, followed by Samtools and Samclip to identify the best matches and store them in a FASTQ file. This allowed to efficiently filter noise in the initial BBDuk output and to calculate precisely the percentage of reads mapping to the full direct repeat in the submitted (+) and reverse (-) orientation. Each archive was analysed with RSeQC (see Methods) and, for stranded libraries, the result was finally adjusted according to the archive orientation to propose a transcription orientation of the input repeat sequence. The TOP workflow is shown in Figure 2.

**Figure 2:**
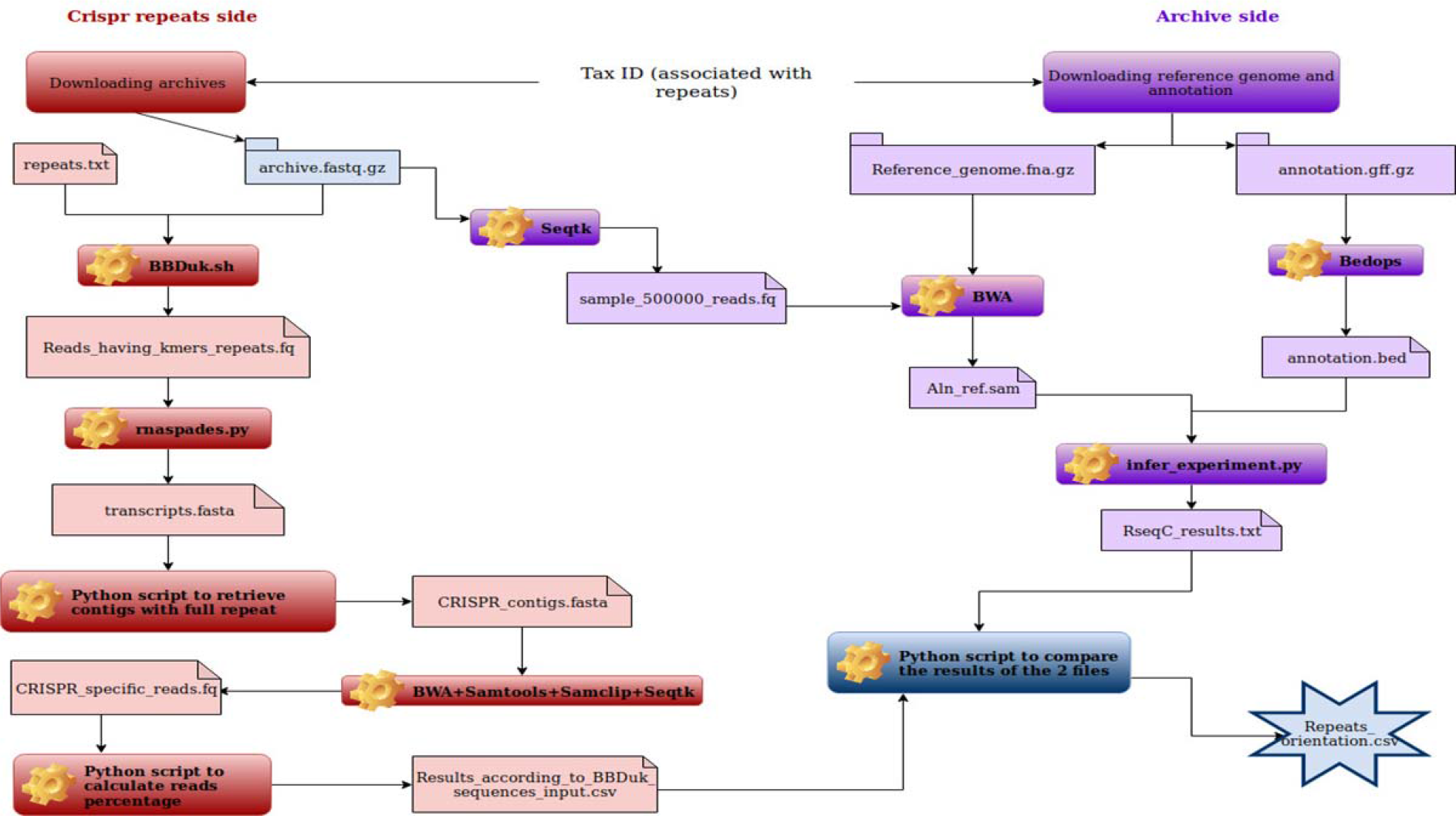
Workflow of the TOP pipeline

### Implementation and validation of TOP

TOP was run with *P. aeruginosa* archive SRR2099777, using 13 different consensus repeats as input, of which eight were variants of the same repeat (in sense and antisense orientation) present in a type I-F system, and five were associated to a type I-E system. The later system could not be investigated because it was not present in the strains used to produce the RNA-Seq archives. For the type I-F repeats, transcription was observed in the selected archive and more than 90% of repeat-specific reads were in the expected orientation.

We then selected the bacterial species *Rhodobacter capsulatus* (taxid 1061) which possessed four el4 CRISPRs with two different repeats (37bp and 32bp long), and for which 18 RNA-Seq archives were available. In CRISPRCasdb, which uses orientation predicted by the CRISPRdirection software (Biswas *et al.*, 2014), the orientation of the 37bp repeats is unknown, and that of 32bp repeat is predicted to be sense. All informative (stranded) archives provided a consistent orientation of the repeats (Table 2), in agreement with the independent prediction made by CRISPRstrand (Alkhnbashi *et al.*, 2014) which appears to be more efficient that CRISPRdirection in this instance.

**Table 1:**
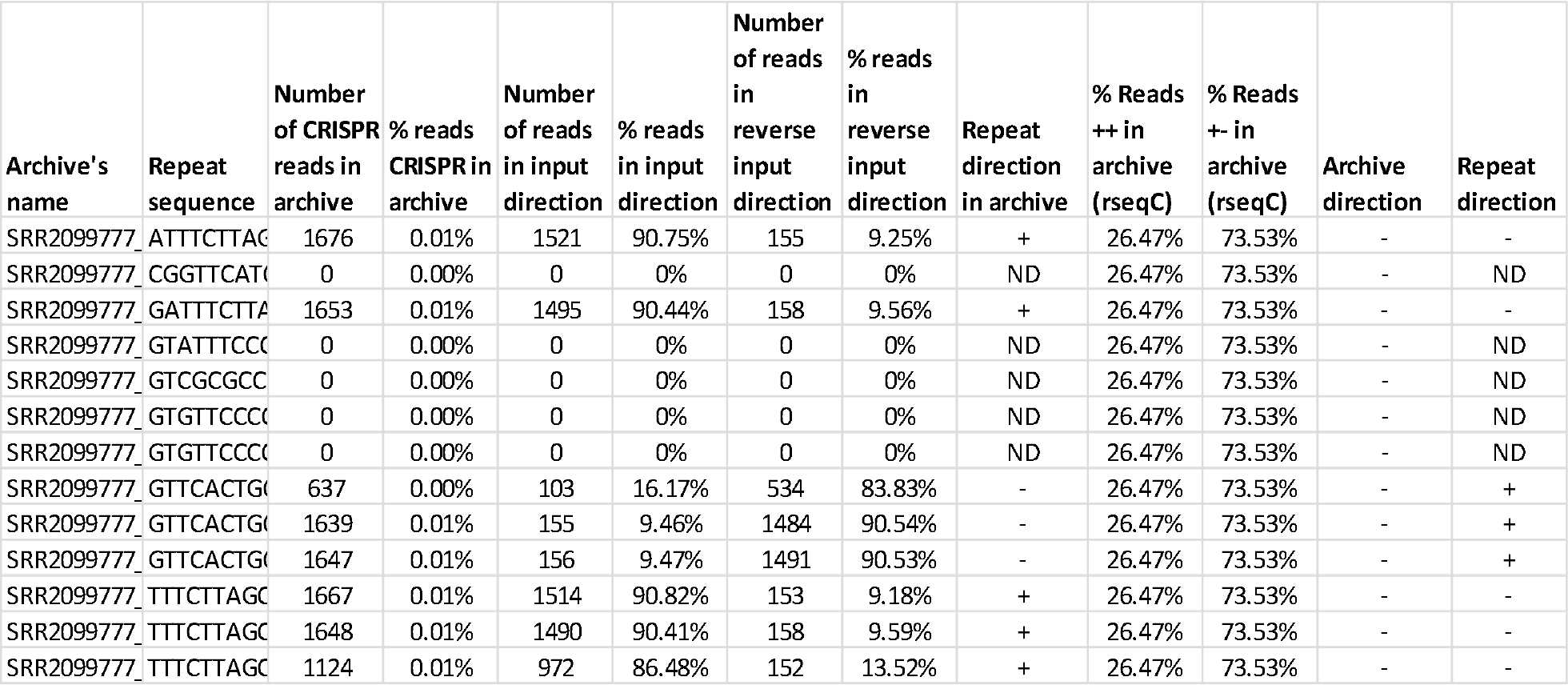
TOP analysis of *P. aeruginosa* RNA-Seq archive SRR2099777 and *X. oryzae* RNA-Seq archive SRR2915666.

**Table 2:**
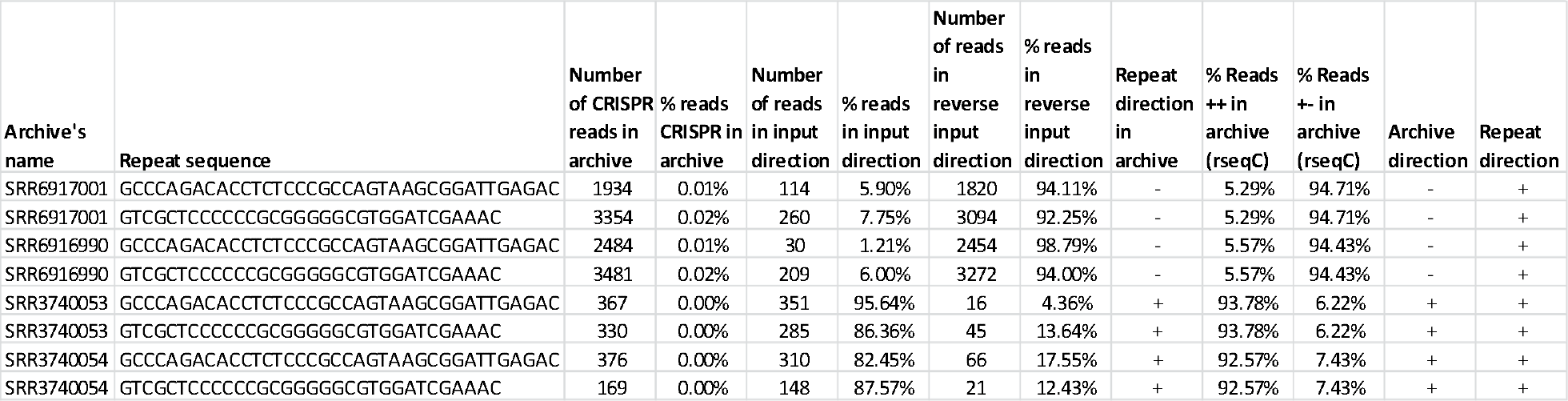
Analysis of four *R. capsulatus* archives with TOP. Two different CRISPR repeat sequences were submitted to the program.

We observed that with some archives TOP did not return any result. With *M. janaschii* archive SRR4240639 TOP failed at the level of contig assembly, possibly due to the large number of CRISPR arrays existing in this genome or to the read quality and amount. Replacing SPAdes with Trinity, another assembly tool, did not improve the result. In the *X. oryzae* archive SRR2915666, 1066 reads were specific for the repeat but the archive was unstranded and therefore of no use.

As a last validation test, we submitted to TOP the sequence of three well-characterized *Staphylococcus aureus* sRNA; Teg23, Teg41 and SAOUHSC_02572 (Carroll *et al.*, 2016). In the three cases correct orientation was obtained showing that the program can be used to analyze sequences other than CRISPR repeats (Table 3).

**Table 3:**
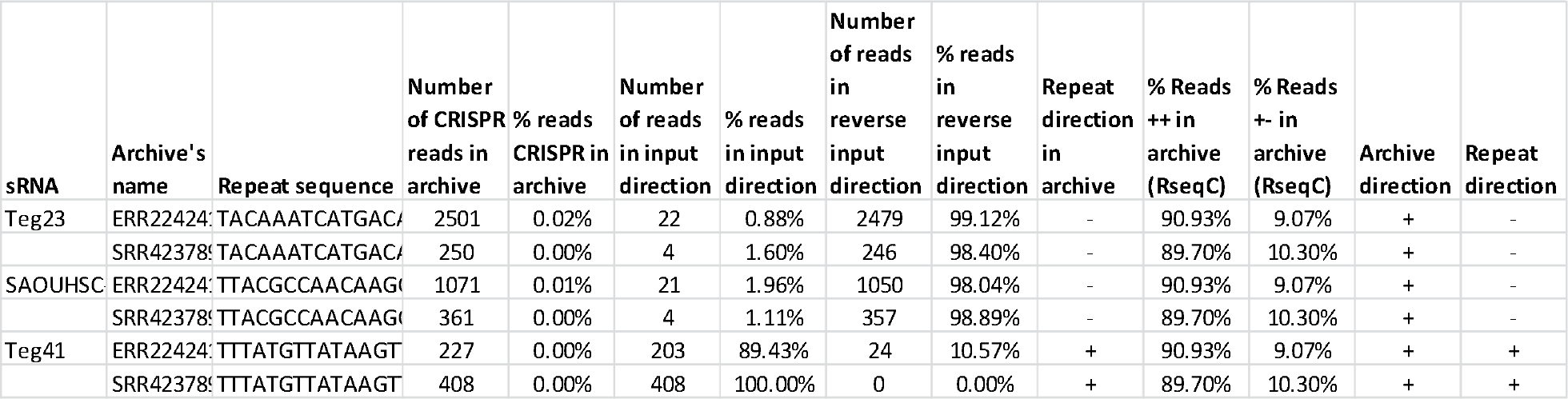
TOP analysis of three *S. aureus* sRNA in two RNA-Seq archives.

### Analysis of *Escherichia coli* data and statistics

To evaluate the number of reads necessary to obtain a reliable result we performed statistical analyses on TOP data from *E. coli* (taxid 562) for which a large number of RNA-Seq archives are available. In this species two CRISPR-Cas systems exist, type I-E and type I-F, present or absent in the available fully sequenced genomes analysed in CRISPRCasdb. A total of 21 repeats derived from the CRISPR arrays can be separated into three groups. One repeat is present in a unique strain (Ecol_422, GCA_002012185.1 assembly accession) on a plasmid devoid of *cas* genes, two repeats (the same sequence in forward and reverse orientation) are present in type I-F CRISPR arrays, and 18 sequences are variants of a repeat present in type I-E CRISPR arrays, in two orientations. Transcription of the CRISPR array in the type I-E system has been highly investigated which makes it a model of choice to test our program (Figure 3) (Pougach *et al.*, 2010).

**Figure 3:**
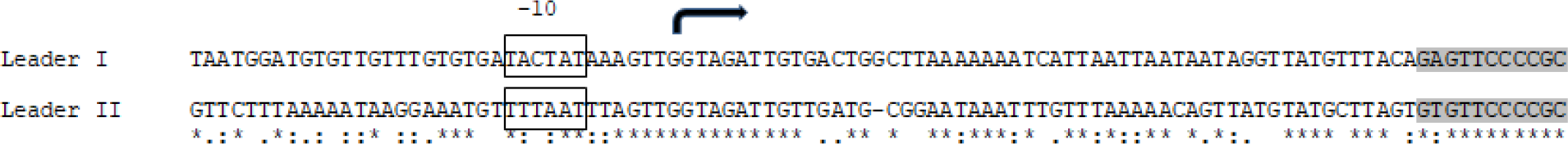
Promoter sequence of *E. coli* type I-E CRISPR array. In the leader preceding two CRISPR arrays, a Pribnow box is present 10 nucleotides before the transcription initiation site shown by an arrow (Pribnow, 1975). The beginning of the first repeat sequence is highlighted in grey.

A total of 521 *E.coli* RNA-seq archives were downloaded from the EBI-ENA repository. TOP analysis of this dataset confirmed the transcription of repeats from the two systems (Table S1 and Table 4). Transcription of type I-E was observed in 174 archives, most of them also containing small amounts of reads for the type I-F CRISPRs. Type I-F reads were present in 53 archives (34 without type I-E) in small amounts and the orientation results were not consistent. No reads were found corresponding to the repeat localized on the plasmid CRISPR array in a single strain. This plasmid was probably not present in the strains used to produce the RNA-Seq archives.

**Table 4:**
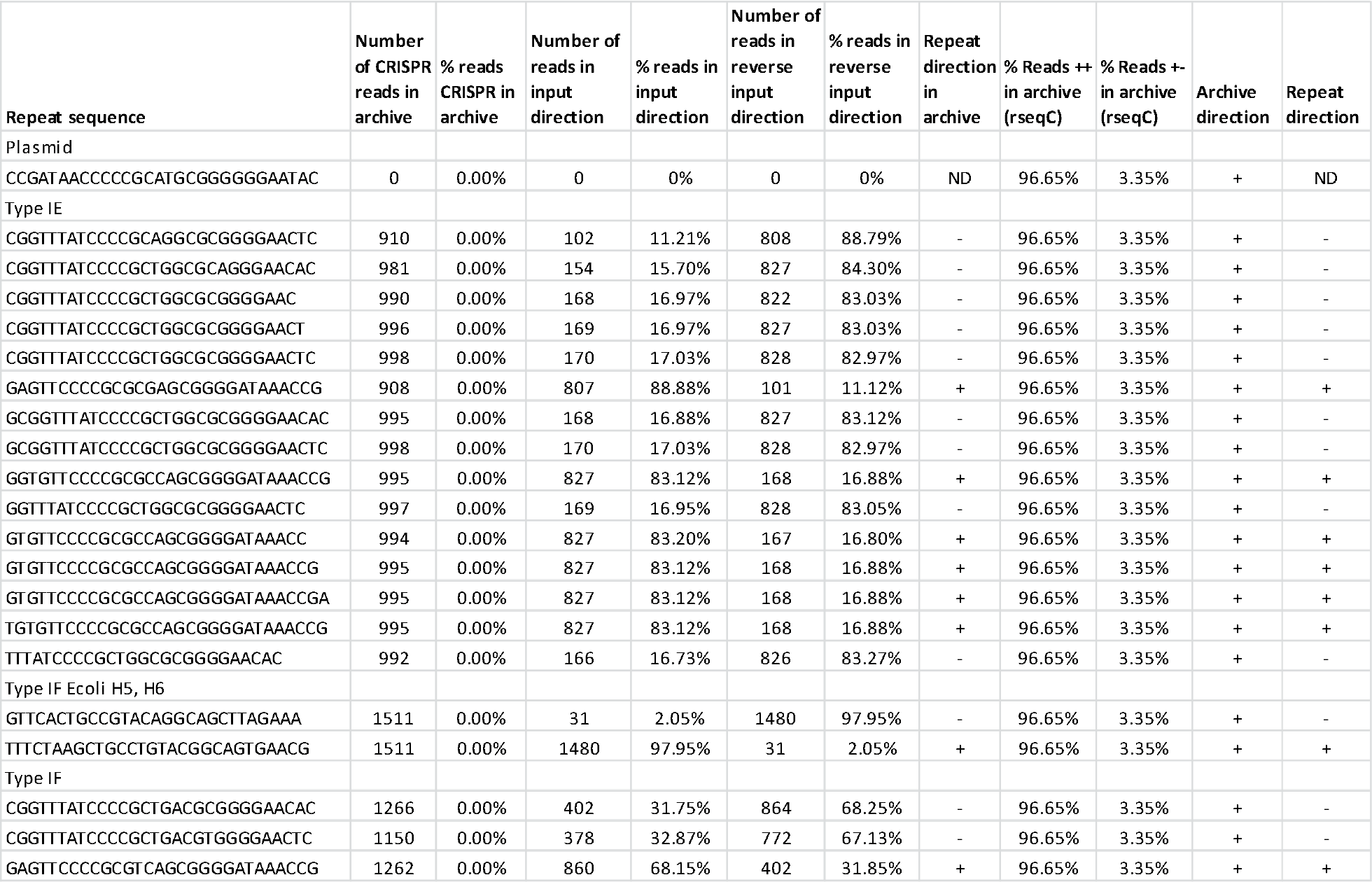
Analysis of archive SRR5051302 from *E.coli* (taxid 562). This archive contains 102 million of 75 bp paired-reads.

Results differed quantitatively between archives but repeat orientations were generally consistent. Several cases of transcription from the two strands in equal amounts were observed. The type I-E system was highly expressed and transcription was similarly oriented in the majority of archives (>= 80%). The same amount of reads was found in the different members of a repeat group, as they are so similar that BBDuk will extract the same reads when using a k-mer value of 19.

It seems obvious that the larger the number of reads, the higher our chance to determine the orientation of the CRISPR. Empirically we observed that the same results were obtained when the number of CRISPR reads aligned to a given element was superior to 100. Another important parameter was the percentage of reads in a given orientation. Indeed in some archives, although the number of CRISPR reads was high, there was not a clear cut difference in terms of transcript level between the two strands. Therefore it appeared that orientation could not always be inferred, even in stranded archives.

We thus performed statistical analyses to test TOP predictions. A binomial test can indicate whether read orientation on a CRISPR locus is random or not. Figure 4 displays the distribution of binomial p-values in function of the number of CRISPRs reads in the different *E. coli* archives. It shows that loci with more than 100 mapped-reads almost always have a significant orientation p-value.

**Figure 4:**
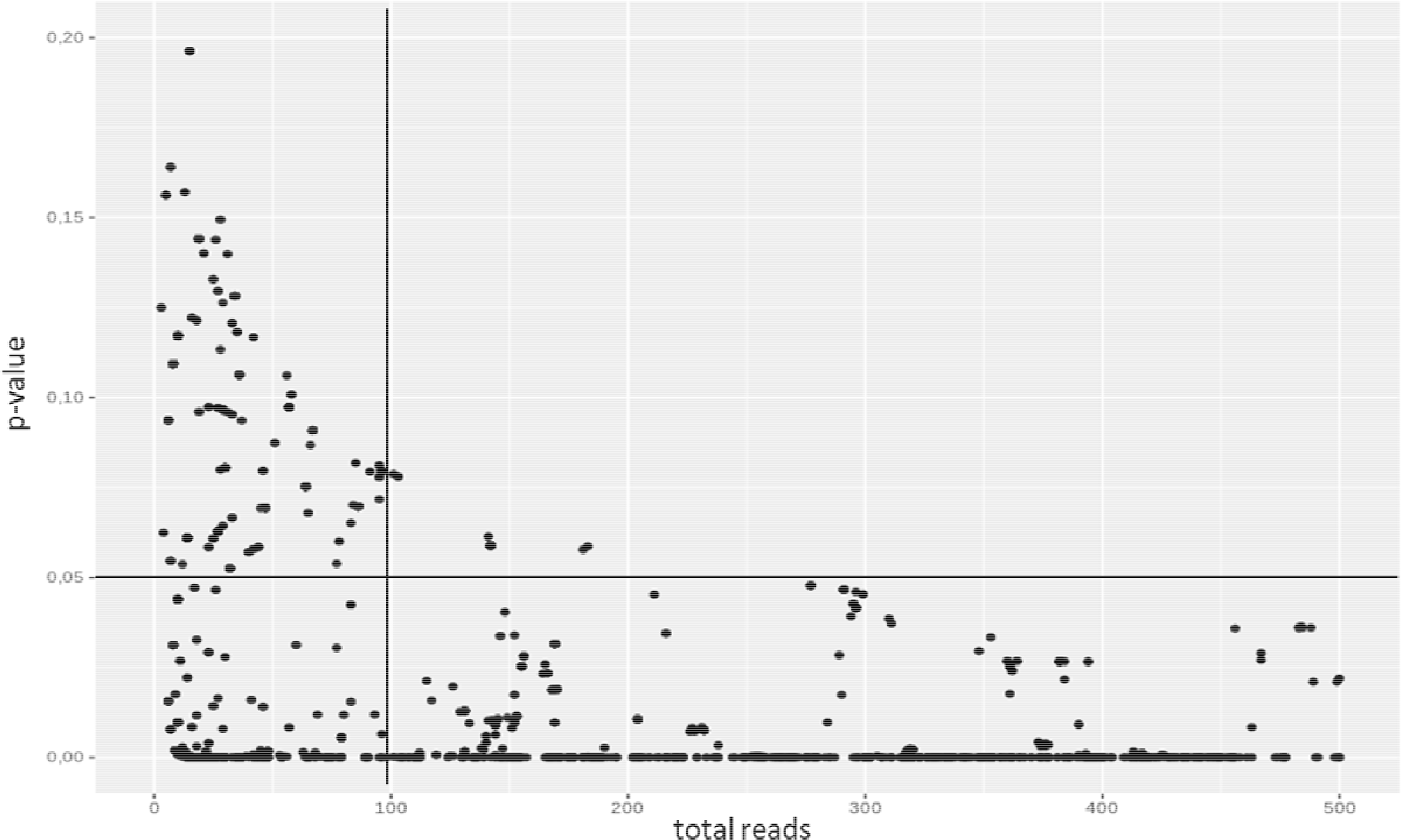
Dot Plot of the p-values according to the numbers of CRISPR reads. For clarity of the figure the total number of CRISPR reads has been limited to 500.

We also generated separated graphs using only stranded archives or unstranded archives to confirm that our data were not biased by the size of the archives (total number of reads) and that, consequently, calling an archive “unstranded” has a biological meaning. Two data sets with 140 archives each were created:

- Stranded archives (true positives) which should produce low p values.
- Unstranded archives (negative control) which should produce higher p values.

The two graphs in Figure 5 A and B show the distribution of the p-values according to the number of reads for both stranded and unstranded archives. It appears that for the stranded archives the p value is low (almost equal to zero), whereas for the unstranded archives the p value is consistently higher than the p value of stranded archives, even if sometimes lower than 5%. Strand bias in unstranded archives is probably due to technical biases in Illumina library preparation and sequencing.

**Figure 5:**
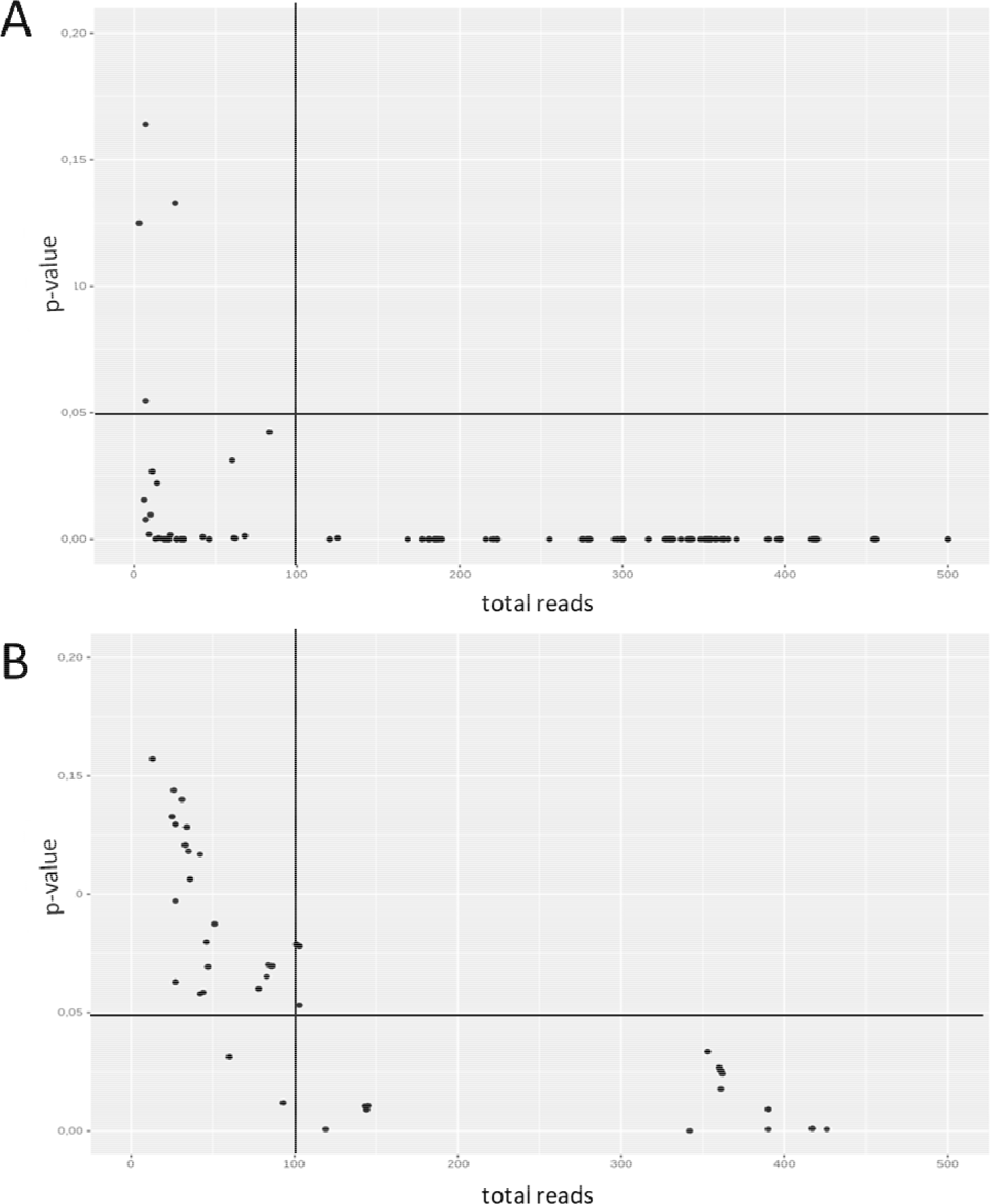
Dot Plot of the p-values according to the numbers of reads for A) 140 stranded archives and for B) 140 unstranded archives.

Regardless of sequencing bias, TOP results can be trusted when the number of CRISPR-specific reads per archive is over 100 or if the p-value is lower than 10^−5^.

### Testing the pipeline on all available archives

The pipeline was run on 4476 RNA-Seq archives for which an el4 CRISPR repeat was identified. A limited number of species accounted for almost half of the available archives: *E. coli* (taxid 562), *S. aureus* (taxid 1280) and *P. aeruginosa* (taxid 287) were represented respectively by 520, 369 and 320 archives while a few others accounted for more than 100 archives such as for example *Streptococcus pyogenes* (taxid 1314), *Clostridioides difficile* (taxid 1496), *M. tuberculosis* (taxid 1773) and *Klebsiella pneumoniae* (taxid 573). A total of 620 el4 consensus repeats were submitted to the program. For 786 archives (17%) the pipeline process failed, whereas no specific transcripts were found in 1996 archives (45%). In 1694 archives (38%) CRISPR reads were present, but part of these archives was unstranded and therefore not useful. Eventually 139 repeats could be analyzed corresponding to 100 different species (Table S2). Table S3 displays the comparison with CRISPRCasdb and CRISPRmap (which makes use of CRISPRstrand) for 73 different repeats in representative archives. TOP provided a result for 53 repeats in 50 different species for which the transcription orientation had not been determined in CRISPRCasdb. For 21 repeats present in 17 species the orientation provided in CRISPRCasdb was not in agreement with TOP and was presumably incorrect. TOP orientated 24 repeats which were not part of the CRISPRmap database accessed December 2019. CRISPRmap and TOP disagreed for ten repeats and agreed for the remaining repeats (53 % of analysed repeats).

Among the discrepancies between TOP and CRISPRmap was the case of *Acinetobacter* sp. ADP1 (taxid 62977) which possesses three CRISPR arrays near a cluster of type I-F *cas* genes. About 80% of CRISPR reads were found in reverse orientation in several independent archives as compared to the prediction of CRISPRmap, and the archive was clearly oriented (97% of the transcription in the sense orientation) (Table 7). The leader was possibly present in the orientation suggested by TOP as revealed by alignment of the CRISPR arrays flanking sequences (not shown). It indicates that some transcription might also be initiated from the other end of the array.

We observed that in some samples there was not a large excess of CRISPR reads in a given orientation in independent stranded archives, suggesting that some transcription was taking place in both directions. It may be explained by the presence of multiple CRISPR arrays in the genome, possibly using different promoters for transcription, such as in *Thermobifida fusca* (taxid 2021) which possesses two CRISPR-Cas systems (type I-E and type III-B) and 12 el4 CRISPR arrays (Table 5).

**Table 5:**
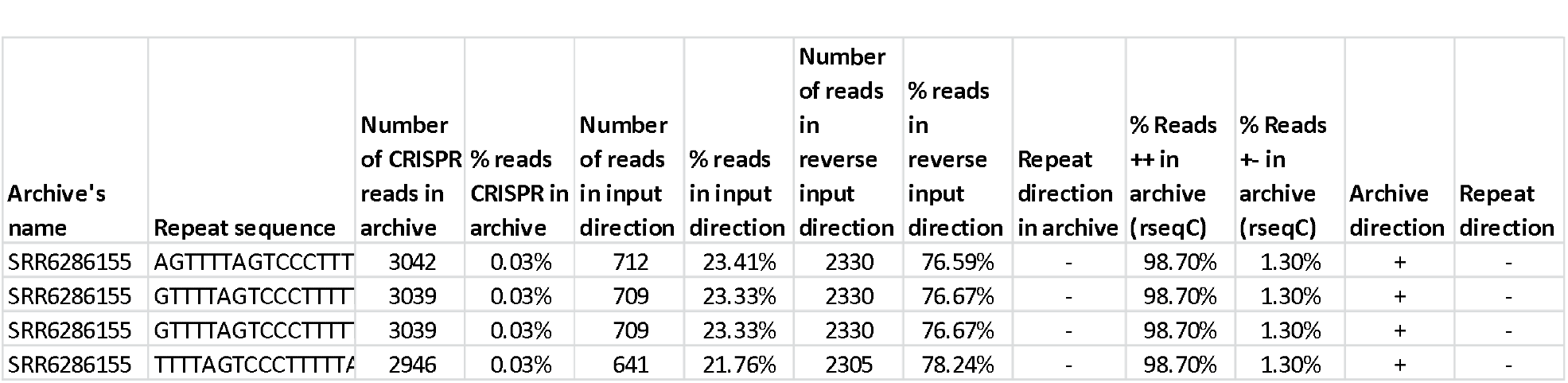
Analysis of two RNA-Seq archives for *T. fusca* (taxid2021).

In *Campylobacter jejuni* (taxid 197) CRISPR reads in sense and antisense direction were respectively around 75% and 25%, although the archives were clearly stranded (Table 6).

**Table 6:**
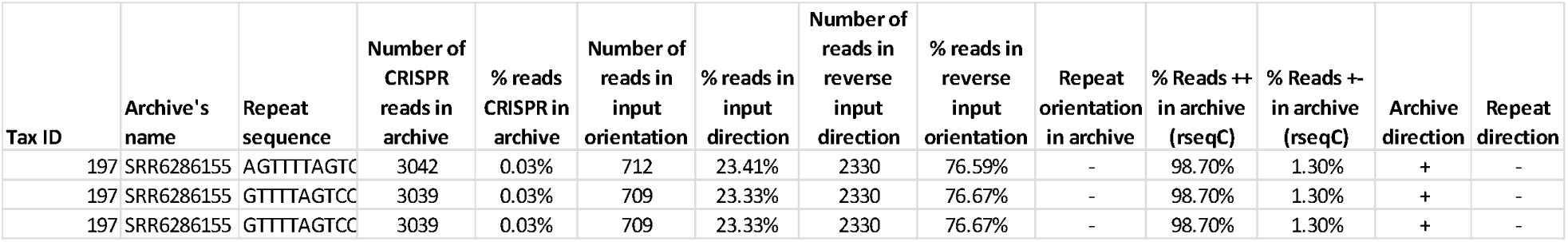
Analysis of *C. jejuni* CRISPR repeats transcription in archive SRR6286155.

**Table 7:**
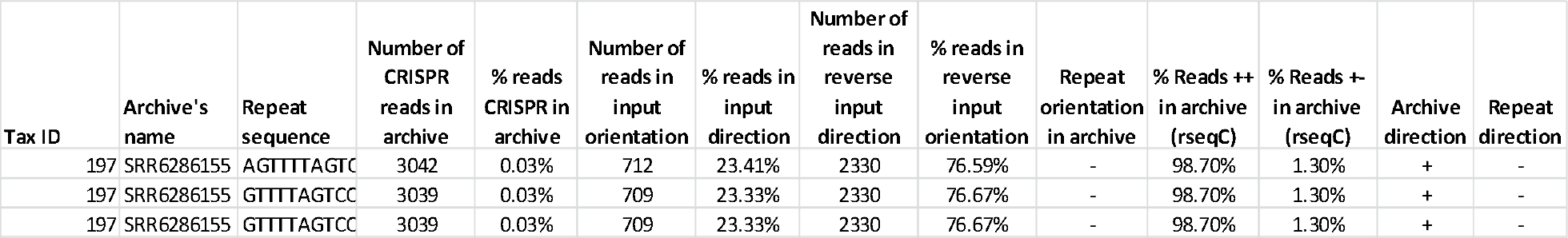
Investigation of *Acinetobacter* ADP1 CRISPR transcription in archive SRR6782867 and SRR6782865.

The AT-rich repeat sequence in *C. jejuni* CRISPR arrays holds a Pribnow box (Figure 6) and it was demonstrated by RNA-seq analysis that transcription was initiated at individual promoters in each repeat in this species and that the transcription was unidirectional (Dugar *et al.*, 2013). It is possible that a second promoter is localized elsewhere, reversely oriented.

**Figure 6:**
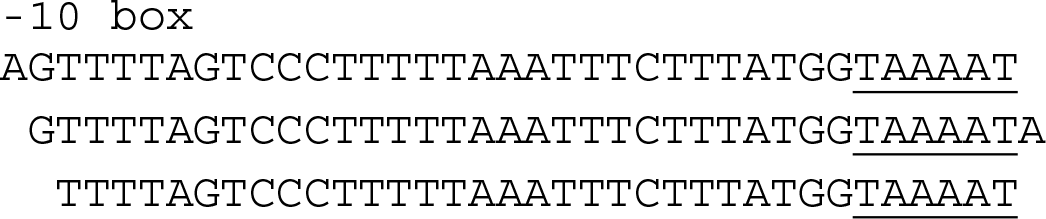
Sequence of the three consensus repeats of *C. jejuni* CRISPR arrays. A Pribnow box is found at the end of the repeat sequence.

### Comparing the level of expression of CRISPR arrays in a genome

TOP analysis of the available RNAseq archives allows comparing the level of expression of different CRISPR arrays within a genome. For example, in *E. coli*, the type I-F array was transcribed 5-10 times less than the type I-E array, and in *Listeria monocytogenes* (taxid 1639), the CRISPR belonging to the type II-A system was transcribed five times more efficiently than the arrays belonging to the type I-B system. A similar observation was made for different *Streptococcus* species (taxid 1308, 1309, 1311) with a type II-A system as compared to type I-C or type I-E systems. Finally high expression was observed for two out of three CRISPRs present on a plasmid in cyanobacteria *Synechocystis* sp. (taxid 1148) and *Synecochoccus* sp. (taxid 32049) associated with type I and type III systems. It rose to 0.65% of the total number of reads in one *Synecochoccus* sp. archive.

## Discussion

The TOP pipeline was used successfully to analyze the transcription orientation of repeats present in CRISPR arrays, as well as other short non-coding sequences using RNA-Seq archives. The capacity to automatically investigate large numbers of archives and sequences allows fast and easy investigation of any sequence of interest. The program can also provide some information on the level of transcription of different sequences in a single genome, although this necessitates that the input sequence maps at a single location in the genome.

Part of these archives was not oriented and for the others it was necessary to identify the orientation. Therefore using RSeQC for orientation estimation was an essential step before investigating the transcription of non-coding sequences.

We observed that the pipeline occasionally failed due to the incapacity of rnaSPAdes to produce a contig with the reads recovered with BBDuk. Further work is necessary to improve this step which is essential to reduce the background of non-specific reads.

Overall the results were in agreement with the predictions of CRISPRmap and provided additional information on repeats that are not part of CRISPRmap database. In some cases it appears that transcription takes place from two different promoters, prompting for additional analyses to better identify these promoters.

## Acknowledgements

We thank Pierre-Albert Charbit for providing information from CRISPRCasdb.

